# Blood flow diverts extracellular vesicles from endothelial degradative compartments to promote angiogenesis

**DOI:** 10.1101/2022.12.19.521008

**Authors:** B. Mary, N. Asokan, K. Jerabkova-Roda, A. Larnicol, I. Busnelli, T. Stemmelen, A. Pichot, A. Molitor, R. Carapito, O. Lefebvre, J.G. Goetz, V. Hyenne

**Author notes:** Equal contribution (J.G. Goetz, V. Hyenne).

## Abstract

Extracellular vesicles released by tumors (tEVs) disseminate via circulatory networks and promote microenvironmental changes in distant organs favoring metastatic seeding. Despite their abundance in the bloodstream, how hemodynamics affect the function of circulating tEVs remains unsolved. We experimentally tuned flow profiles in vitro (microfluidics) and in vivo (zebrafish) and demonstrated that efficient uptake of tEVs occurs in endothelial cells subjected to capillary-like hemodynamics. Such flow profiles partially reroute internalized tEVs towards non-acidic and non-degradative Rab14-positive endosomes, at the expense of lysosomes, suggesting that endothelial mechanosensing diverts tEVs from degradation. Subsequently, tEVs promote the expression of pro-angiogenic transcription factors in flow-stimulated endothelial cells and favor vessel sprouting in zebrafish. Altogether, we demonstrate that capillary-like flow profiles potentiate the pro-tumoral function of circulating tEVs by promoting their uptake and rerouting their trafficking. We propose that tEVs contribute to pre-metastatic niche formation by exploiting endothelial mechanosensing in specific vascular regions with permissive hemodynamics.

## Introduction

Inter-organ communication is instrumental in maintaining systemic homeostasis, in coordinating metabolic response to environmental challenges or in reacting to diseased tissue. It can be mediated by hormones, cytokines, but also by membrane-covered structures belonging to the heterogenous family of extracellular vesicles (EVs) (Tkach and Théry, 2016). These small vesicles (30nm-5μm diameter range) transport bioactive molecules (RNAs, lipids, proteins) between distant organs by traveling through circulatory networks (Yáñez-Mó et al., 2015; Cheng and Hill, 2022). In cancer, as the disease becomes systemic and progresses from primary to secondary sites, EVs actively contribute to organ cross-talk and thereby impact both primary tumor growth and metastatic spreading. Tumor EVs (tEVs) released by primary tumors spread via blood or lymphatic circulation and reach distant organs where they alter the microenvironment (Kalluri and LeBleu, 2020; Marar et al., 2021). When reaching future metastatic organs before tumor cells arrival, tEVs modify their cellular and extracellular composition and create a pre-metastatic niche favorable to metastasis formation (Peinado et al., 2017; Ghoroghi et al., 2021a). While the changes induced by tEVs on distant organs started to be unraveled over the past years, the transit of tEVs from the circulation to the new organ are very poorly described. Notably, while circulating tEVs are mostly taken up by endothelial cells (together with intravascular patrolling monocytes) in both vascular blood (Morishita et al., 2015; Imai et al., 2015; Takahashi et al., 2013; Hyenne et al., 2019) and lymphatic settings (García-Silva et al., 2021), few studies described their fate in a realistic vascular environment (van Niel et al., 2022; Verweij et al., 2019b; Hyenne et al., 2019). By contrast, the behavior of circulating tumor cells in the vasculature is better understood (Follain et al., 2020). For instance, it is now well established that blood flow forces directly impact the arrest, extravasation and metastatic capacities of circulating tumor cells (Follain et al., 2020). We thus hypothesize that hemodynamics are likely to tune the vascular targeting and uptake of tEVs and thereby control how they prime distant organs and form pre-metastatic niches. Furthermore, hemodynamics strongly affects the biology of the endothelium: through a variety of mechano-sensors, endothelial cells sense and rapidly respond to flow forces, notably to shear stress, by adapting their architecture and homeostasis (Fang et al., 2019). For instance, hemodynamic forces alter endothelial shape and cytoskeleton organization (Li et al., 2005), but also endosomal pathways, such as endothelial autophagy (Vion et al., 2017). Flow shear forces also directly impact the capacity of endothelial cells to take up material flowing in the circulation, such as ions, proteins or nanoparticles, although it is not clear whether they alter their fate (Han et al., 2015, 2012; Tarbell, 2010). Therefore, we speculated that hemodynamics would impact the fate and trafficking of internalized tEVs. To test this possibility, we built an experimental pipeline that allows to control flow forces while assessing the initial steps of tEVs targeting, uptake and fate in realistic hemodynamic environments. We combined an in vivo model, the zebrafish embryo, that we adapted to the study of tEVs (Hyenne et al., 2019; Verweij et al., 2019a), together with an in vitro microfluidics models where a monolayer of endothelial cells can be challenged with tunable flow profiles (Follain et al., 2018). Such multi-modal approach allowed to document, for the first time, how hemodynamics control the trafficking of tEVs upon uptake, whose consequences are instrumental in mediating their microenvironmental priming function. We show that hemodynamics not only promote the uptake of tEVs, they also further regulate their fate and function through the control of their trafficking routes in endothelial cells. Importantly, we show that blood flow promotes a partial lysosomal escape which enhances the pro-angiogenic function of tEVs.

## Results and Discussion

### Blood flow tunes endothelial uptake of circulating tEVs

To determine the influence of hemodynamics on the dissemination of circulating tEVs, we took advantage of the zebrafish embryo that we previously described (Hyenne et al., 2019). At 48h post-fertilization, the zebrafish embryo is composed of a complex vascular network suited to real-time imaging of circulating EVs (Verweij et al., 2021, 2019a; Hyenne et al., 2019). When injected in the Duct of Cuvier, injected tEVs quickly disseminate through the bloodstream and reach the caudal plexus via the dorsal aorta (Fig.1a and (Hyenne et al., 2019)). While the dorsal aorta carries circulating tEVs at relatively high flow velocities (Average > 550 μm/s), these quickly return in venous compartments characterized by low flow velocities (Average < 450 μm/s) (Follain et al., 2018). Interestingly, when assessing which vascular regions were preferably targeted by tEVs, we observed that both endothelial cells and patrolling macrophages of low flow regions efficiently internalized circulating tEVs (Hyenne et al., 2019). We first validated tEVs uptake by endothelial cells by correlating intravital imaging with electron microscopy (iCLEM) (Fig. 1a, iCLEM panel). We then carefully and concomitantly measured blood flow velocities and endothelial accumulation of circulating tEVs (Fig.1a). To do so, we isolated tEVs from tumor cells by differential centrifugation, labeled them with MemGlow (Hyenne et al., 2019) and injected them in the circulation of zebrafish embryos (Mary et al., 2020). We analyzed four vascular regions (Fig.1a, hemodynamics panel) that are representatives of the flow profiles found in the caudal plexus and plotted the amount of tEVs internalized by endothelial cells with respect to the flow velocities (Fig.1a and Movie 1). We observed a significant inverse correlation between flow profiles and endothelial accumulation of tEVs (Pearson correlation coefficient r=-0,65; p=0,021): while accumulation of tEVs is maximal in venous regions with 400 to 450 μm/s flow velocity, it is significantly reduced in arterial regions where flow ranges from 575 to 650 μm/s. In order to confirm that flow profiles directly control the uptake of tEVs, we experimentally and pharmacologically tuned blood flow velocities in the ZF embryo and quantified the resulting tEVs uptake in a common vascular region (caudal vein). Decreasing flow velocities with lidocaine (Fig.1b, left), a sodium channel blocker that reduces pacemaker activity of the heart (Follain et al., 2018), lead to a significant decrease of tEVs uptake by endothelial cells (Fig.1b, right). Conversely, increasing the heart pacemaker activity (Fig.1c, left) using IBMX (3-isobutyl-1-methylxanthine) (Follain et al., 2018) results in a slight albeit non-significant increase in tEVs uptake in the venous endothelium (Fig.1c, right). To confirm these in vivo data suggesting that tEVs uptake is mostly efficient at lower velocity flow profiles of 400 μm/s, we exploited in vitro microfluidics system, which allows for a precise and homogenous control of flow regimes (Fig.1d, left). Similarly, we measured the internalization of fluorescent tEVs perfused on a monolayer of endothelial cells and observed that tEVs accumulate more efficiently in endothelial cells subjected to a 400 μm/s flow speed when compared to static (or no flow) conditions (Fig.1d, right). Altogether, our in vitro and in vivo experiments demonstrate that endothelial cells most efficiently accumulate tEVs when subjected to flow velocities of 400 μm/s. We propose that binding and arrest of tEVs at the endothelial surface are prevented when flow velocities exceed the permissive value in the range of 400 μm/s while both uptake and trafficking of tEVs would be impaired, through mechanosensing, at low flow velocities. Interestingly, we had observed EVs rolling on the surface of the endothelium of zebrafish embryos at this flow velocity (Hyenne et al., 2019). In addition a velocity of 400 μm/s favors the arrest of circulating tumor cells (Follain et al., 2018) and is in the range of flow speed measured in small capillaries from organs that are prone to metastasis such as human liver or mouse brains (Follain et al., 2020). Nevertheless, hemodynamics cannot be the sole explanation for such behavior and we and others had shown that endothelial targeting of tEVs also relies on specific adhesion receptors present at their surface, such as integrins or CD146/MCAM (Hoshino et al., 2015; Ghoroghi et al., 2021b; Jerabkova-Roda et al., 2022). Therefore, efficient binding of tEVs to the endothelial surface occurs when adhesion-prone tEVs exploit permissive flow profiles. Interestingly, similar observations were made with synthetic nanoparticles as their uptake by endothelial cells is higher at venous-like low shear stress than in static conditions, but decreases at higher arterial-like shear stress in a receptor-dependent manner (Han et al., 2012, 2015; Chen et al., 2020; Lin et al., 2010). Therefore, is it likely that the destination of circulating tEVs relies on the combined effect of hemodynamics and receptor-ligand interactions.

**Fig. 1.**
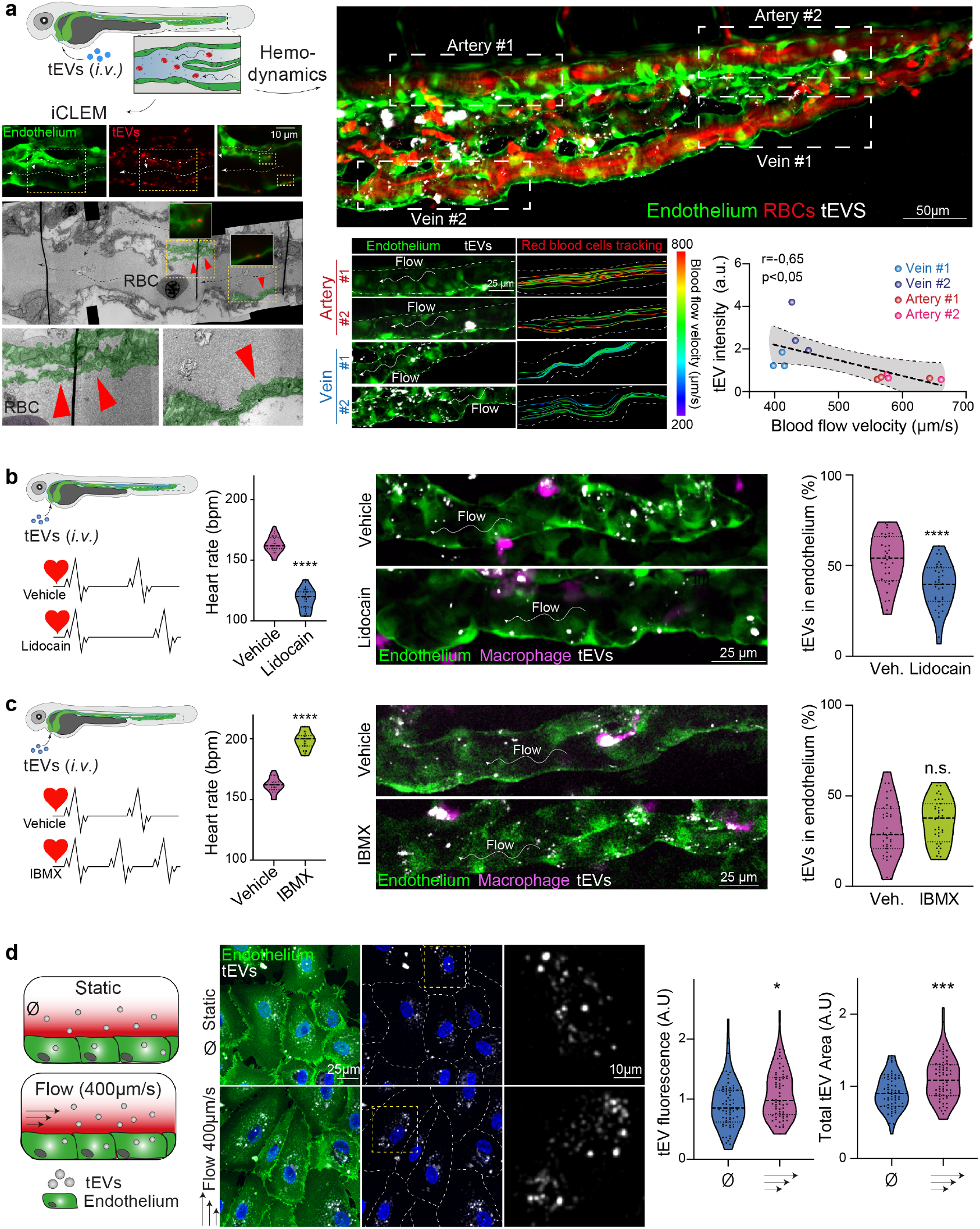
Blood flow tune the uptake of circulating tEVs by endothelial cells a) Description of the experimental setup: 2 days post-fertilization (Fli1:GFP (endothelium) Gata1:dsRed (red blood cells -RBCs) zebrafish embryos are injected intravascularly with Memglow-Cy5-labeled 4T1 tEVs and imaged in the caudal plexus after 30 minutes (right, Z projection). Velocities of individual circulating RBCs are tracked and fluorescent tEV signal internalized in the endothelium is measured in four indicated regions per fish (2 aortic and 2 veinous). Bottom left: representative images of the four regions showing tEVs within the endothelium (Z projections) and RBCs tracks with a color code representing velocities. Bottom right: graph showing the correlation between internalized tEVs and RBC velocities in the same region (n= 3 embryos; Pearson r=-0,65; p=0,021). b-c) Memglow-Cy5-labeled 4T1 tEVs are injected intravascularly in Fli1:GFP (GFP endothelium) embryos with decreased (b) or increased (c) heart pacemaker activity with respectively lidocaine and IBMX treatments. Left graphs display pacemaker activities (1 dot: 1 fish, p<0,0001 Student T-test). Middle: representative images, maximum intensity projections. Right graph represents the percentage of fluorescent tEVs within the endothelium (1 dot: 1 fish; Lidocain: n=36 per condition, three independent experiments; p<0,0001 Student T-test; IBMX: n=39 per condition, three independent experiments, p=0,11 Student T-test). d) Memglow-Cy5-labeled 4T1 tEVs are perfused on HUVECs endothelial monolayer labeled with Memglow 488 in static or flow conditions (400 μm/s) and imaged by confocal microscope after 3h. Representative images and quantification of internalized tEVs showing increase in tEV total area (p=0,0001 Mann Whitney), in tEV total fluorescence (p=0,013 Student T-test) (data per cell, normalized to static; 1 dot: 1 field of view; n=67; experiment performed in quintuplicate).

### Circulating tEVs are partially re-routed to alternative non-acidic and non-degradative RAB14 positive compartments

We then aimed to characterize the fate of circulating tEVs in flow-stimulated endothelial cells. To investigate the trafficking routes exploited by tEVs, we generated vHUVECs endothelial cell lines stably expressing fluorescent markers of early (mCherry-Rab5), late (mEmerald-Rab7) and recycling endosomes (eGFP-Rab11) as well as late endosome-lysosomes (RFP-LAMP1). We tracked the internalization of fluorescently labelled tEVs by live imaging and quantified their colocalization with endosomal markers after 3h using an automated image analysis pipeline (see Material and Methods). We found that internalized tEVs mostly colocalize with LAMP1 and, to a lesser extent with Rab5 and Rab7 positive compartments (Fig.2a and Fig.S1a-d). Flow had no effect on the distribution of tEVs among these endo-somal markers suggesting that endothelial mechanosensing had no impact on the general internalization routes used by tEVs. The accumulation of circulating tEVs in LAMP1 positive compartments suggests that they could mostly be stored in degradative compartments, similarly to what happens in patrolling macrophages (Hyenne et al., 2019). Therefore, we interrogated whether flow could divert tEVs from degradative machineries. We first labeled degradative compartments in endothelial cells using the Magic Red membrane permeant probe, which detects the activity of the lysosomal protease cathepsin B. While most of the internalized tEVs accumulate in Magic Red positive compartments, flow significantly reduced the proportion of tEVs found in cathepsin B-positive structures (Fig.2b) suggesting that internalized tEVs are partially redirected to non-degradative compartments upon flow mechanosensing. Since the degradative activity of lysosomes hydrolases relies on the acidic luminal pH (Perera and Zoncu, 2016), we next probed the pH of tEV-containing compartments by exploiting the pH-sensitive reporter pHluorin. When anchored on EVs external membrane via an insertion in the extracellular domain of the tetraspanin CD63, such construct, which is also fused to pH-insensitive mScarlet at its intracellular C-terminal end (Sung et al., 2020), allows to probe the pH of the compartments targeted by tEVs (Fig.2c, left). We generated tumor cells expressing CD63-pHluorin-Scarlet and validated that these cells can secrete pH-sensitive fluorescent EVs (data not shown). The ratio of green (pHluorin) towards red (mScarlet) fluorescence was then used to probe the pH of tEVs’ environment. When pH-sensor tEVs were perfused on endothelial cells, we again observed striking differences in endothelial cells subjected to flow. We found that flow drastically increased the pHluorin/mScarlet ratio (Fig.2c) suggesting that endothelial mechanosensing can change the trafficking routes of internalized tEVs and reroute them towards more neutral compartments. Together, these results show that internalized tEVs are in part diverted to less acidic and less degradative compartments when endothelial cells are cultured under flow. In order to identify such compartments, we focused on Rab14, a Rab GTPase known for being localized in LAMP1 positive late endosomes, in addition to other compartments (Hoffman et al., 2022). Among other functions, Rab14 is involved in intracellular virus and pathogen trafficking towards late endolysosomes (Kyei et al., 2006; Okai et al., 2015; Kuijl et al., 2013). Importantly, Rab14 also controls the transit of internalized material toward non-acidic LAMP1 positive compartments (Trofimenko et al., 2021). Therefore, we stably expressed GFP-Rab14 in endothelial cells and first confirmed that Rab14 is mostly absent from lysosomal degradative compartments, as it only weakly co-localizes with Magic Red in static and flow conditions (Fig.S2). We then assessed whether tEVs traffic through Rab14 compartments. While internalized tEVs are often found in close proximity to Rab14 positive independently of flow, the proportion of tEVs within the lumen of Rab14 positive compartments is significatively increased in flow-stimulated endothelial cells (Fig.2d). Altogether, our results show that the moderate flow regimes partially switch EVs trafficking toward Rab14 positive non-acidic and non-degradative compartments suggesting that endothelial mechanosensing of flow would prevent tEVs from degradation. To validate our observations in vivo, we injected CD63-pHluorin-mScarlet tEVs in the circulation of zebrafish embryos where the flow speed was pharmacologically manipulated. We measured the pHluorin/mScarlet ratio of tEVs internalized in a single vessel (caudal vein) as performed in Fig.1 and observed that the pHluorin/mScarlet ratio was reduced in embryos with decreased flow velocities (Fig.2e), suggesting that tEVs accumulate in more acidic compartments under low flow velocity. Conversely, increasing blood flow velocity with IBMX increases the pHluorin/mScarlet ratio, suggesting that tEVs tend to accumulate in less acidic compartments in endothelial cells facing higher flow speed. Altogether, our combined in vitro and in vivo data show that circulating tEVs follow a different trafficking route depending on hemodynamic forces applied on endothelial cells. At moderate or high velocities, internalized tEVs are partially rerouted towards non-acidic compartments suggesting that the fate of tEVs, and their function, is strictly controlled by the mechano-sensing abilities of the endothelium. Flow-dependent trafficking of internalized tEVs could determine the proportion of EVs undergoing lysosomal degradation and tEVs allowed to transfer their cargo through back-fusion in endosomes. However, since both processes rely on endosomal acidification (Bonsergent et al., 2021; Joshi et al., 2020), the extent to which tEVs escape lysosomal degradation could relate to alternative fate, such as direct signaling from endosomes (Shelke et al., 2019) or endothelial transcytosis which allows tEVs to cross the blood-brain barrier (Morad et al., 2019).

**Fig. 2.**
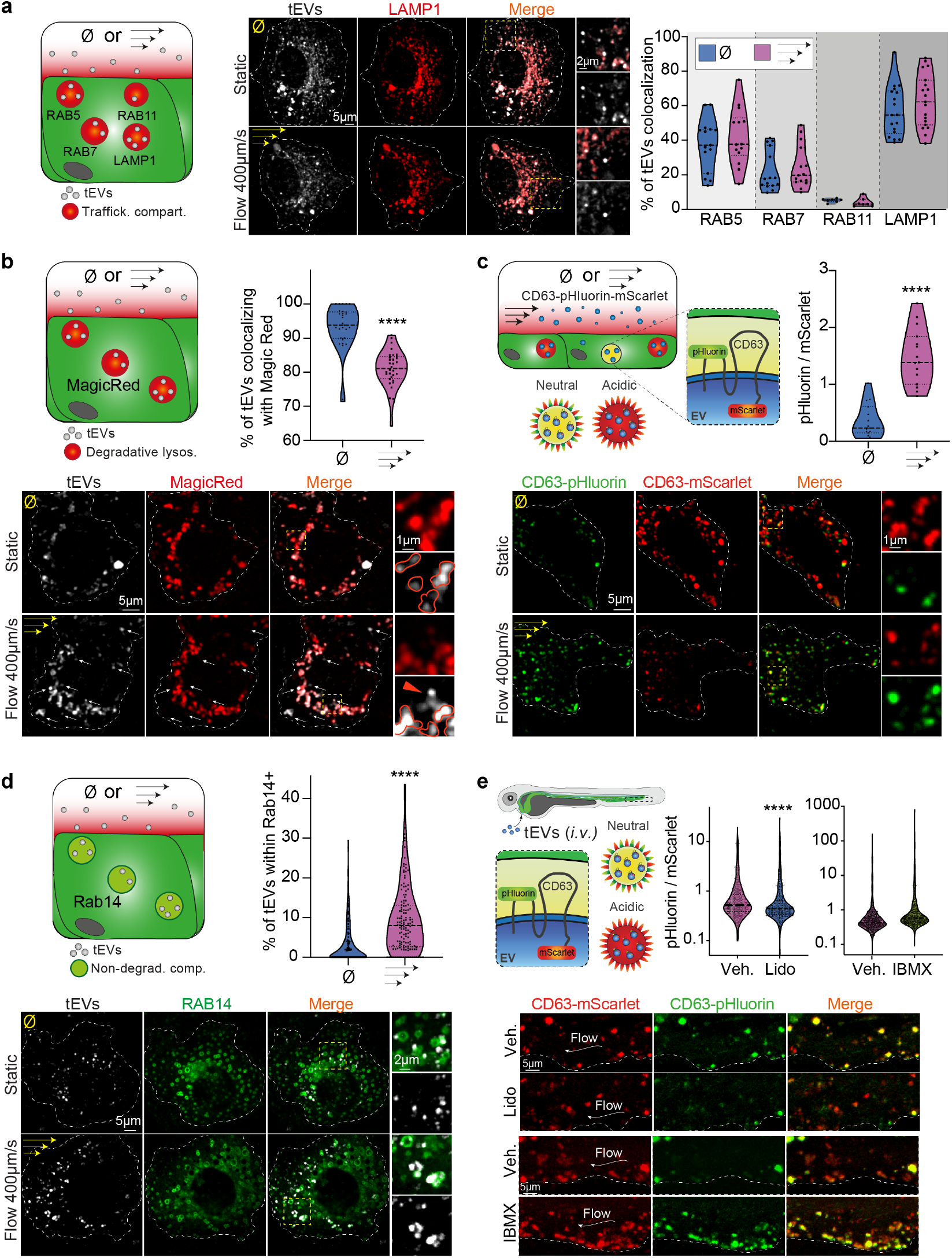
Blood flow diverts tEVs sorting towards RAB14 positive non-degradative and non-acidic compartments. a) In vitro, internalized tEVs co-localize more efficiently with LAMP1 than RAB5, RAB7 and RAB11 in endothelial cells cultured under flow. Memglow-Cy5 labeled tEVs were perfused on vHUVECs cells stably expressing FP-RAB5,7,11 or LAMP1 cultured in flow or static conditions and imaged by spinning disk after 3h (single plane representative images). Automated colocalization analysis (Mann Whitney, one dot= one cell, n=7-20). b) Decreased accumulation of circulating tEVs in degradative compartments in endothelial cells cultured under flow. Memglow-Cy5 labeled tEVs were perfused on HUVECs cultured in flow or static conditions and labeled using Magic Red to stain compartments with cathepsin B activity. Representative spinning disk single plane images (Manual quantification, one dot=one cell, n=30, Mann Whitney, p<0,0001). c) tEVs accumulate in less acidic compartments when endothelial cells are cultured under flow. CD63-pHluorin-mScarlet 4T1 EVs were perfused on HUVECs cultured in flow or static condition and imaged by spinning disk after 3h (single plane representative images). Graph represents the pHluorin/mScarlet ratio (one dot = one field of view, n=15, experiment performed in triplicate Mann Whitney, p<0,0001). d) tEVs show increased accumulation in Rab14 positive compartments in endothelial cells cultured under flow. Memglow-Cy5 labeled tEVs were perfused on vHUVECs cells stably expressing GFP-RAB14 cultured in flow or static conditions and imaged by spinning disk after 3h (single plane representative images). Manual and automated quantifications show increased colocalization in flow conditions (one dot=one cell, n>139, Mann Whitney, p<0,0001). e) In zebrafish embryos, modulating flow speed alters tEVs trafficking in acidic compartments. CD63-pHluorin-mScarlet 4T1 tEVs were injected in wild-type zebrafish embryos treated with lidocaine or IBMX to respectively decrease or increase heart beat rates and flow velocity. Maximum projection of fish caudal vein area showing tEVs accumulation in the endothelium (visualized by transmitted light and represented with a white dashed line) 3h post-injection. Graph represents the pHluorin/mScarlet ratio in endothelial cells (one dot = one Z plane, n>500, at least 15 embryos from three independent experiments, Mann Whitney, p<0,0001).

### Blood flow promotes lysosomal pathways in endothelial cells

Endothelial cells have exceptional mechanosensing abilities that orchestrate multiple cellular functions (Fang et al., 2019; Freund et al., 2012). Having observed that flow re-directs tEVs towards non-degradative compartments, we wondered whether this resulted from a broad re-organization of endosomal trafficking in flow-stimulated endothelial cells. To address this question, we simply analyzed endolysosomal trafficking in flow-stimulated endothelial cells, in absence of tEVs. We first compared transcriptional response using RNAseq in endothelial cells subjected to 400 μm/s or not (Table 1). Interestingly, such flow profiles significantly favor the transcription of genes associated with lysosomal pathways (Fig.3a). When probing the endolysosomal pathway using fluorescent dyes, we observed that flow increased the number of lysotracker positive compartments (Fig.3b). When carefully assessing inner trafficking compartments using electron microscopy, we confirmed that flow significantly increases the number of endolysosomes, which displayed a smaller mean size (Fig.3c) in agreement with fluorescence microscopy (Fig.3b), and that is directly linked to their degradative capacities (de Araujo et al., 2020). Therefore, to gain insight into the functionality of those compartments, we quantified the cathepsin B activity using the Magic Red dye. We observed that the number of Magic Red positive compartment is increased (Fig.3d). This result suggests that endothelial cells sensing flow profiles that are permissive to tEVs uptake also increase their lysosomal degradative activity, which is regulated by pH. Altogether, these experiments show that flow profiles that do favor tEVs uptake and re-routing also promote endolysosomal degradative pathways. Therefore, the re-routing of EVs toward non-degradative compartments does not solely result from a mechano-transduction pathway that adjusts lysosomal pathways, but more likely reflects a flow-dependent EV specific switch in trafficking route. Interestingly, the RNAseq analysis further revealed that flow-stimulated endothelial cells activate expression of genes involved in the regulation of lysosomal pH, including the proton exchanger NHE9 (SLC9A9) which limits endosome luminal acidification (Fig.3a)(Kondapalli et al., 2015; Beydoun et al., 2017). We confirmed by independent RNA sequencing: that flow triggers endogenous NHE9 expression and generated endothelial cells stably expressing NHE9-mCherry. Us-: ing these, we observed that fluorescent tEVs accumulate in NHE9-positive compartments (Fig. 3e). Therefore, flow not only strengthens the lysosomal pathway in endothelial cells, it also triggers the formation of NHE9 compartments where EVs accumulate. Importantly, by increasing endosomal pH, NHE9 limits the lysosomal degradation of activated receptors such as the EGFR and enhances their signaling activity (Kondapalli et al., 2015). Therefore, it is tempting to speculate that the flow-mediated alternate trafficking of tEVs in less acidic compartments impacts their function.

**Fig. 3.**
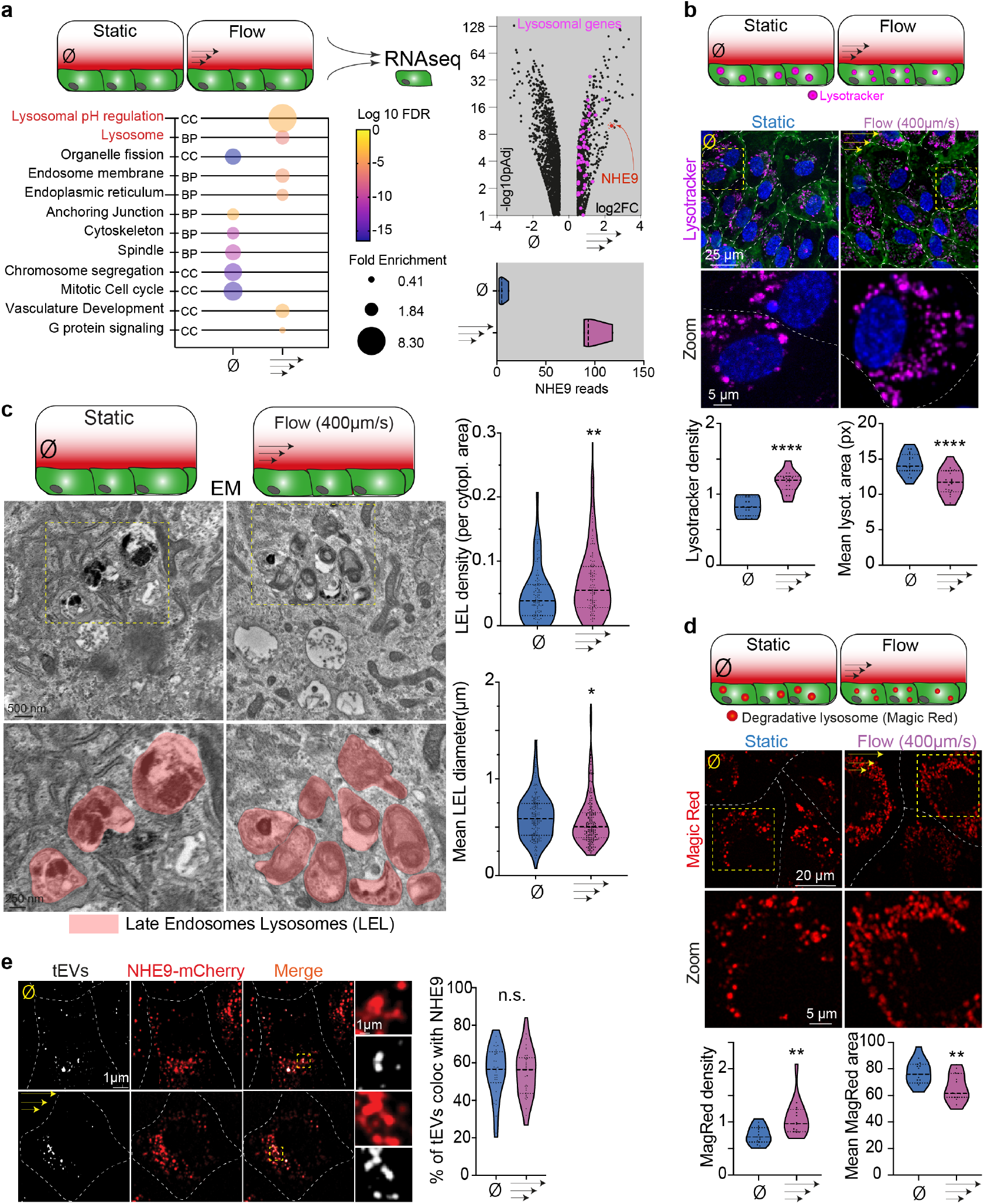
Blood flow upregulates lysosomal pathway. a) RNA sequencing of HUVEC cells cultured under a moderate flow speed reveals an enrichment in genes associated with lysosomal regulation. Transcriptomics was performed on HUVEC cells cultured in static or flow conditions. The bubble plot shows the GO terms differentially expressed between static and flow conditions (CC: Cellular components; BP: biological process). The volcano plot shows the genes differentially expressed between static and flow conditions highlighting the 73 genes associated with lysosomes in pink and NHE9 in red. The lower right graph shows the number of NHE9 reads in static and flow conditions. b-c) Increased number of lysosomes in endothelial cells cultured under flow and observed by photonic (b) and electronic microscopy (c). HUVEC cells were cultured in static or flow conditions, labeled with Memglow 488, Nucblue and lysotracker to visualize cells, nuclei and lysosomes respectively and imaged by confocal as shown on representative maximum projection images. Left graph represents the relative number of lysosomes per cell and right graph the mean area per lysosome (one dot is one field of view, n=20, p<0,0001, Student T-test) c) Representative electron microscopy images of HUVEC cells cultured in flow and static conditions and schematic representation of late endosome lysosome (LEL, red). Graphs show an increase in the number of LEL per cytoplasmic surface (one dot represents one field of view; n=121 and 125 fields of view respectively; p=0,0028 Mann Whitney) and a decrease in their average diameter in HUVEC cells cultured under flow (one dot represents one LEL; n=166 and 178 LELs respectively; p=0,015 Mann Whitney). d) Increase in degradative compartments in HUVEC cells upon flow treatment. HUVEC cells were cultured in static or flow conditions, labeled Magic Red to visualize compartments with cathepsin B activity and imaged by confocal as shown on representative maximum projection images. Left graph represents the relative number of magic red positive compartment per cell (1 dot= 1 field of view (FOV); p=0,0025 T-test) and right graph the mean area of these compartments. (1 dot = 1 FOV p=0,049 Mann Withney). e) NHE9 positive compartments accumulate tEVs. Representative confocal images (single plane) of internalized EVs in vHUVECs cells expressing NHE9-mCherry in flow or static conditions. Colocalization was quantified using an automated pipeline. (Each dot represents one field of view, n=34; two independent experiments, Mann Whitney, p=0.6417).

**Table 1:** List of genes differentially expressed between HUVEC cells cultured in flow (positive fold change) and static conditions.

### Blood flow and circulating tEVs cooperate to favor angiogenesis

Having demonstrated that blood flow favors both the uptake and the lysosomal escape of tEVs, we wondered whether they might tune endothelial response. We first interrogated to what extent blood flow and tEVs would cooperate and impact transcriptional programs of targeted endothelial cells. Interestingly, the endothelial transcriptome was differentially impacted by tEVs in static and flow condition (Fig.4a and Table 2). While the number of genes dysregulated by tEVs is similar in static and flow, their identity differs: genes whose transcription was impacted when tEVs were internalized by flow-stimulated endothelial cells remain unaltered when tEVs were internalized in static endothelial cells (Fig.4a, right). This demonstrates that endothelial cells respond differently to tEVs when subjected to flow. As tEVs escape lysosomal compartments in such conditions, this suggests that such endosomal re-routing allows tEVs to transfer their message that ultimately tunes gene expression. Whether this transfer results from direct signaling from endosome membranes or from cytoplasmic cargo release remains to be deciphered. Among the genes upregulated by tEVs in flow-stimulated endothelial cells, we found several pro-angiogenic transcription factors, such as ID1, ID2, ID3, Hey1, Hey2, MAFB, Runx1 and HES1 (Benezra et al., 2001; Fischer et al., 2004; Morioka et al., 2014; Kitagawa et al., 2013). We further identified genes that are involved in two pro-angiogenic signaling pathways, Notch (HEY1, HEY2, HES1, JAG1) and TGFß (Smad6, Smad7, Bambi, PMEPA1, Nog) (Fig.4a, right). Mechanistically, both pathways can be activated directly from endosomes (Baron, 2012), and eventually from internalized tEVs (Shelke et al., 2019). How tEVs activate these pathways remains unclear, yet we found that they contain several regulators of the TGFß pathway (Smad5, Smurf2, etc.) (Ghoroghi et al., 2021b), including TGFß type II receptor, whose presence on tEVs is sufficient to trigger the pathway in receiving cells (Xie et al., 2022). Altogether, we demonstrate here that blood flow and tEVs cooperate to favor lysosomal escape allowing tEVs to promote a pronounced pro-angiogeneic transcriptional program in endothelial cells. Activation of this program, potentially through Notch and TGFß pathways, could impact metastasis in many ways. Activation of Notch1 in endothelial cells of the premetastatic niche, for instance, favors neutrophils infiltration and ultimately metastasis (Wieland et al., 2017). Besides, endothelial tEVs were shown to activate the Notch pathway in endothelial cells, resulting in increased expression of HEY1 and HEY2 and formation of capillary-like structures in vitro and in vivo (Sheldon et al., 2010). In addition, activation of the TGFß pathway in endothelial cells promotes inflammation and endothelial permeabilization (Chen et al., 2019), while the presence of TGFß type II receptor on tEVs correlates with metastasis (Xie et al., 2022). While pro-angiogenic programs, which could possibly be activated by the Notch and TGFß pathways, require a concerted action of flow and tEVs uptake in endothelial cells, whether they impact the angiogenic activity of endothelial cells remained unsolved. To test the relevance of such gene signature in vivo, we adapted a well-established experimental tumor angiogenesis assay in zebrafish embryos (Nicoli and Presta, 2007) to investigate whether tEVs could impact endothelial response in realistic hemodynamic conditions. To this end, we assessed the ability of intravenous injected and circulating tEVs to promote the formation of neo-sprouts from existing sub-intestinal vessels in embryos bearing a tumor mass. When embryos are injected with tEVs, the tumor-induced sprouting is potentiated with an increased number of neo-vascular sprouts per embryo (Fig.4b). Importantly, tEVs also increase the percentage of embryos bearing tumor-induced endothelial sprouts when compared to embryos injected with PBS. These results confirm that circulating tEVs promote tumor-induced neo-angiogenesis in vivo, a scenario that is likely to happen when tEVs shape pre-metastatic niches (Peinado et al., 2017; Ghoroghi et al., 2021a). While tumor tEVs were previously shown to promote angiogenesis in vitro (Todorova et al., 2017), our results suggest that such effect can be potentiated by hemodynamic forces of perfused vessels. As a consequence, we expect circulating tEVs to favor the formation of neo-vascular sprouts in capillary-like vessels that are prone, from a hemodynamics stand-point, to favor arrest of CTCs (Follain et al., 2018). The formation of new and abnormal blood vessels could constitute a first step in the creation of pre-metastatic niches, leading to the subsequent recruitment of specific immune populations and ultimately favoring homing of circulating tumor cells (Peinado et al., 2017) (Fig. 4c). We have recently shown that endothelial cells also hijack a flow-stimulated pro-angiogenic transcriptional program to perform intravascular remodeling that favors the extravasation of arrested CTCs (Follain et al., 2018, 2021). It is tempting to speculate that tEVs could mediate such intravascular remodeling, whose dependence on flow forces is also established (Follain et al., 2021). Finally, one could expect that tEVs are continuously secreted after metastasis initiation, leading to the induction of neo-sprouts which sustain the growth of metastatic foci that have colonized distant organs (Kienast et al., 2010)(Fig. 4c). Overall, our work demonstrates for the first time that the fate and function of tEVs flowing in the bloodstream and on their way to shape pre-metastatic niches is tightly linked to hemodynamic forces as well as mechanosensing abilities of the endothelium. We provide here the first evidence that flow-sensing, by diverting internalized tEVs from lysosomal degradation, changes their signaling capacities. We further identified permissive flow regimes where tEVs uptake by endothelial cells is optimal. Such regimes not only favor the uptake and lysosomal escape of tEVs, they also allow the arrest of CTCs that precedes metastatic extravasation and outgrowth (Follain et al., 2020, 2018). Therefore, EVs uptake and CTCs extravasation are likely to occur at similar locations, implying that CTCs would be prone to reach vessels that were already corrupted by tEVs. tEVs could activate endothelial cells locally to favor arrest and extravasation of CTCs and immune cells to further support metastatic outgrowth. In endothelial cells subjected to such flow regimes, internalized tEVs are partially re-routed to non-acidic and non-degradative RAB14 and NHE9 positive compartments with direct and functional consequences on their angiogenic potential. In conclusion, hemodynamics potentiate the functional impact of tEVs on endothelial cells towards a pro-angiogenic response which ultimately shapes pre-metastatic niches and metastatic outgrowth.

**Fig. 4.**
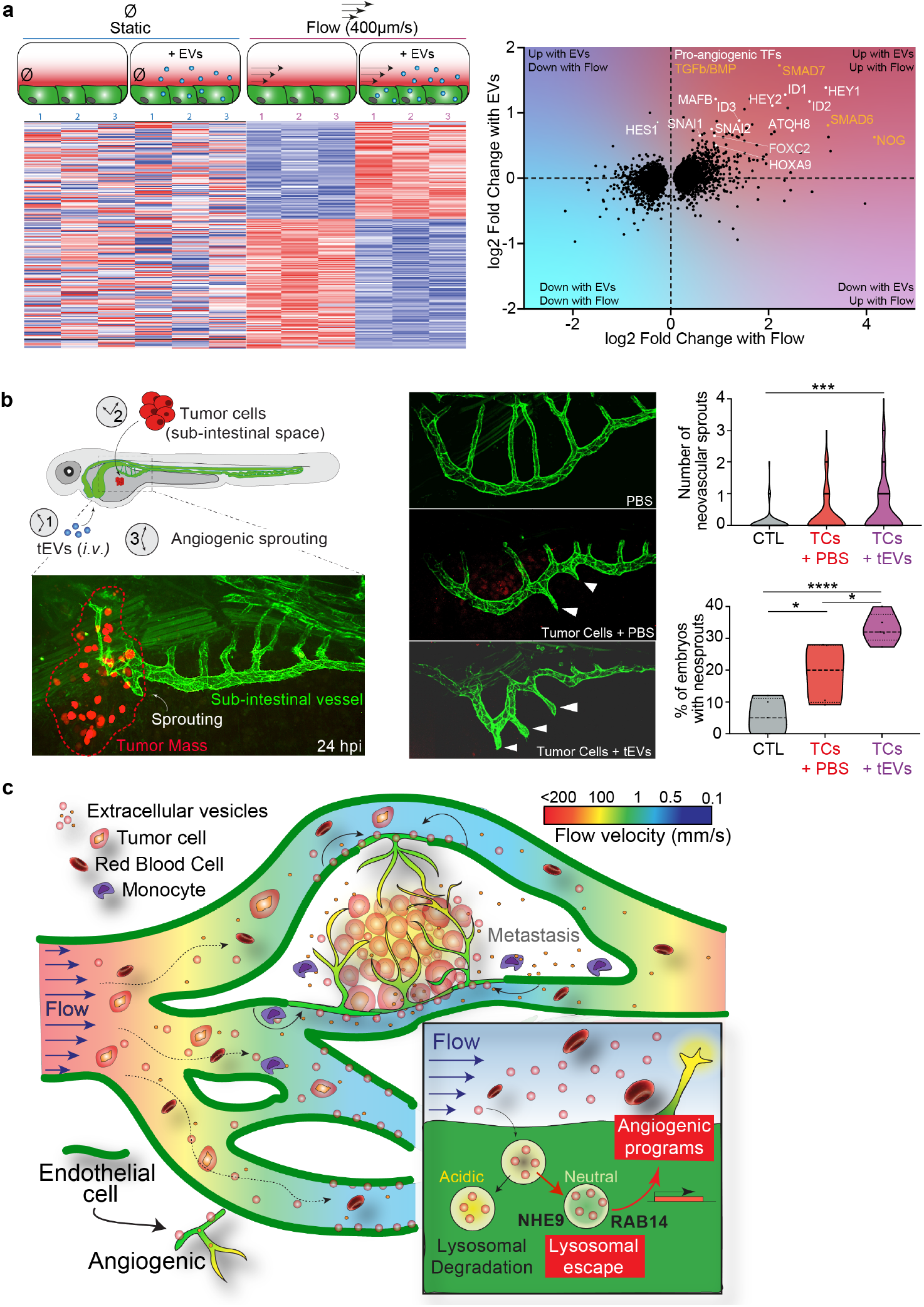
Circulating tumor EVs and blood flow cooperate to promote angiogenesis a) RNA sequencing reveals an enrichment in pro-angiogenic pathways in HUVEC cells cultured under a moderate flow speed and treated with 4T1 tEVs. Transcriptomics was performed on HUVEC cells cultured in static or flow conditions and treated with 4T1 tEVs or PBS for 24h. Heat map shows that the genes differentially expressed upon tEV treatment in flow condition are not deregulated in static conditions. The right graph shows differentially expressed genes with flow and tEVs (pAdj<0,1). Pro-angiogenic transcription factors (orange) and members of the TGFß /BMP pathway (green) are highlighted. b) Tumor EVs promote angiogenesis in vivo. Zebrafish embryos were injected with 4T1 tEVs or PBS and subsequently with 4T1 tumor cells (TCs). Representative confocal images show neovascular sprouts in the sub-intestinal vessels (SIV) 24h post-tumor cell injection. The number of neovascular sprouts (upper graph; one dot=one fish; n 80 embryos from 5 independent experiments; one-way anova with Dunn’s multiple comparison test, adjusted p value=0.0001) and the percentage of embryos with neovascular sprouts (lower graph; one dot=one experiment; one-way anova with Tukey’s multiple comparisons; adjusted p values were as follows: CTL vs PBS+TCs p=0.0186, CTL vs tEVs+TCs p<0.0001 and PBS+TCs vs tEVs+TCs p=0.0156) were quantified. c) model explaining the fate and function of circulating tumor EVs: low flow speed promotes EVs uptake by endothelial cells followed by partial lysosomal escape and rerouting in RAB14 and NHE9 positive compartments. As a consequence, tumor EVs induce a pro-angiogenic response.

**Table 2:** List of genes differentially expressed between HUVEC cells cultured in flow (positive fold change) and static conditions and treated with tEVs (positive fold change) or PBS.

## Methods

### Cell culture

4T1 cells were cultured in RPMI-1640 medium complemented with 10per cent Foetal Bovine Serum (FBS, Hyclone) and 1per cent penicillin/streptomycin (PS, 100U/ml, Gibco). HUVEC cells (Human Umbilical Vein Endothelial Cell, Promocell) were grown in Endothelial Growth Medium (ECGM, Promocell) complemented with supplemental mix (SupplementMix, Promocell) and 1per cent PS. VeravecTM HUVECs human endothelial cells (Angiocrine Biosciences), referred here as vHUVECs, were cultured in ECGM MV2 (Promocell) complemented with 20per cent of FBS and 1per cent PS. For all the microfluidic experiments, HUVECs or vHUVECs were used before passage number 4. HEK293T cells were grown in DMEM (Gibco) complemented with 10per cent FBS and 1per cent PS. All cell lines were maintained at 37C and 5per cent CO2 and verified for the absence of mycoplasma by PCR on a regular basis.

### Stable cell line generation

Lentivirus containing the following constructs were produced in HEK293T cells using JetPRIME (Polyplus, FRANCE) transfection: pLSFFV-mCherry-Rab5A, pLSFFV-RFP-LAMP1, pLSFFV-mEmerald-Rab7A, pLSFFV-eGFP-Rab11A, pLSFFV-GFP-Rab14 (gift from N. Vitale, INCI, Strasbourg, France), pLSFFV-NHE9-mCherry (gift from K. C. Kondapalli, Michigan-Dearborn University, USA), pLSFFV-pHluorin-CD63-mScarlet (gift from A. M. Weaver, Vanderbilt University, USA). vHuvec or 4T1 cells were infected by lentivirus in the presence of 5μg/mL polybrene (Sigma) followed by antibiotic selection (Puromycin at 1μg/mL or blasticidin at 5μg/mL).

### EVs isolation and labelling

4T1 EVs were previously characterized by electron microscopy, nanoparticle tracking analysis, proteomics and western-blots (Hyenne et al., 2015; Ghoroghi et al., 2021b). In brief, cells were cultured at sub-confluency in EV depleted medium for 24h. EV depleted medium was prepared using 2X complete medium, centrifuged at 100,000g for 20h (Optima XE-90 ultracentrifuge - Beckman Coulter) to eliminate EVs from FBS. Supernatant was then filtered at 0,22μm (Millipore) and adjusted to 1X. tEVs were isolated using differential ultracentrifugation protocols as described previously (Hyenne et al., 2015, 2019). The pellets obtained at the final step of 100,000g centrifugation were resuspended in sterile PBS1X. When needed, tEVs were labeled using 200nM of MemGlow-Cy5 (Cytoskeleton Inc.) lipidic dye between the 1st and the 2nd 100 000g ultracentrifugation as described previously. Following the tEV isolation, their number and size distribution were measured by ZetaView NTA (PMX-120-12B-R2 – Particle Metrix). tEVs were stored at 4°C and used within 24h.

### Microfluidics experiments

For microfluidic experiments, IBIDI^®^ μ-slides with 1 (μ-Slide I 0.4 Luer ibiTreat: 1.5 polymer coverslip) or 6 (μ-Slide VI 0.4 ibiTreat: 1.5 polymer coverslip) channels were coated with fibronectin (SIGMA) for 1 hour and seeded with 100 000 or 50 000 cells per channel respectively. Once endothelial cells reached confluency, the channels were either maintained in static conditions or continuously perfused under a flow of 400μm.s-1 for 17h using a Reglo Digital MS-CA 2/12 peristaltic pump. tEVs were then either applied on the static cells or perfused continuously under flow at a concentration of 5.108 to 1.109 particles/ml. After 3h of tEV perfusion, channels were removed from the flow, and washed 3 times with fresh ECGM and 3 times with ECGM/Hoechst 33342 (NucBlue™ ThermoFisher). After 10min incubation, cells were washed twice with ECGM/HEPES 20mM before imaging. To assess lysosome number or Cathespin-B activity, endothelial cells were incubated after EV perfusion with Lysotracker Deep Red (Thermo Fisher Scientific) diluted at 50nM or MagicRed dye (Bio-Rad) diluted at 1:260 respectively for 1h. When needed, cells were incubated with MemGlow-488 (Cytoskeleton Inc.) at 200nM to label the plasma membrane.

### Photonic microscopy and image analysis

Cells were imaged live at 37°C with 5percent CO2 with an Olympus Spinning Disk (60X objective, N.A. 1.2), with a Leica SP5 confocal (63X objective, N.A. 1.25) or with a Zeiss LSM 800 confocal (63X objective, N.A. 1,4). Zebrafish embryos were imaged live at 28°C with a Lecia SP5 confocal equipped with a HC PL APO 20X/0,7 IMM objective or with an Olympus Spinning Disk equipped with a 30X objective (N.A. 1.05). Image analysis and processing were performed using the Fiji (2.0.0) (Schindelin et al., 2012), Cell Profiler (4.2.1) (Stirling et al., 2021) and ICY (2.4.2.0.) (De Chaumont et al., 2012) software. Number, intensity and area of compartments (Fig. 1d, 3b and 3d) were quantified on maximum intensity projections using the ICY Spot detector plugin. To quantify tEV signal in the endothelium in vivo (Fig. 1b and 1c), we used an in-house built Fiji macro to quantify the total tEV fluorescence in the tail region and specifically in the endothelium. Colocalization (Fig. 2a, b, d, Fig. 3e and Fig. S2) was analyzed on processed single Z planes (Despeckle and Gaussian blur filter in Fiji) based on intensity thresholding using Cell Profiler. Areas of overlapping regions between tEVs and given compartment were quantified. The percentage of tEV colocalization with the given compartment was calculated as a ratio of tEV overlap area to the total area of tEV objects times 100. The in vitro CD63-pHluorin/mScarlet experiments (Fig. 2c), were analysed on processed single planes, using masks to detect single spots and measure fluorescence intensities in the endothelium. For each field of view, total green and red spots fluorescence was then measured and used to calculate a pHluorin/mScarlet ratio. For the in vivo CD63-pHluorin/mScarlet experiments (Fig. 2c), the venous endothelium was first manually segmented and the total green and red spots fluorescence in the endothelium was then measured on tresholded single Z planes and used to calculate a pHluorin/mScarlet ratio.

### Electron microscopy

Chemical fixation: Cells were fixed with 2,5 per cent glutaraldehyde (GA)/2,0 per cent paraformaldehyde (PFA) (Electron Microscopy Sciences) in 0.1M NaCac buffer (pH 7.4) at room temperature for 2h, then rinsed in 0.1M NaCac buffer (pH 7.4) (Electron Microscopy Sciences) and post-fixed with 1 per cent OsO4 (Electron Microscopy Sciences) and 0.8 per cent K3Fe(CN)6 (Sigma-Aldrich) for 1h at room temperature. Then, samples were rinsed in 0.1M NaCac buffer (pH 7.4) followed by a distilled water rinse and stained with 1 per cent uranyl acetate, overnight at 4°C sheltered from the light. The samples were stepwise dehydrated in Ethanol (50 per cent x10min, 70 per cent x10min, 95 per cent x15min and 100 per cent 3×10min), infiltrated in a graded series of Epon (Ethanol 100 per cent/Epon 3/1, 1/1) 1h and kept in Ethanol 100per cent/Epon 1/3 overnight at room temperature. The following day, samples were placed in pure Epon 3 x1h and polymerized at 60°C 48h. 100 nm thin sections were collected in 200 copper mesh grids and data set was acquired with a TEM Hitachi 7500 TEM, with 80 kV beam voltage, and the 8-bit images were obtained with a Hamamatsu camera C4742-51-12NR. The number of late endosome-lysosomes per surface of cytoplasm were quantified using the Fiji software.

### Intravascular injection of tEVs in zebrafish embryos

Zebrafish embryos were obtained from the following strains: Tg(fli1a:eGFP), Tg(Fli:LA-eGFP) Tg(Fli1:Gal4; UAS: RFP), Casper Tg(Gata1:RFP; flk:GFP). Embryos were maintained at 28°C in Danieau 0.3X medium, supplemented with 1-Phenyl-2-thiourea (PTU, Sigma-Aldrich) after 24 h post fertilization (hpf). All injection experiments were carried out at 48 hpf and imaged between 48 hpf and 72 hpf. All animal procedures were performed in accordance with French and European Union animal welfare guidelines and supervised by local ethics committee (Animal facility A6748233; APAFIS 2018092515234191). At 48 hpf zebrafish (ZF) embryos were dechorionated and mounted on a 0.8per cent low melting agarose pad containing 650 μM tricaine (ethyl-3-aminobenzoate-methanesulfonate). Embryos were injected intravascularly in the duct of Cuvier with 27,6 nL of Membright Cy5-labeled tEVs or 4T1CD63pHluorinmScarlet-tEVs (at 10E10 EVs/ml) with a Nanoject microinjector 2 (Drummond) under a M205 FA stereomicroscope (Leica) as described previously (Mary et al., 2020; Hyenne et al., 2019).

### Sample preparation for Correlative Light and Electronic Microscopy of ZF embryos

Correlative Light and Electron Microscopy was performed as previously described(Hyenne et al., 2019; Mary et al., 2020). Transgenic fli1:GFP embryos were injected with Memglow-Cy3 tEVs and imaged alive with a Leica SP8 confocal microscope. Z stack was performed on the caudal vein regions that had internalized EVs. After imaging, the embryo was chemically fixed with 2,5per cent glutaraldehyde and 4per cent paraformaldehyde in 0.1 M Cacodylate buffer (the fish tail was cut off in the fixative). The sample was kept in fixative at RT for 1-2h and stored in fixative at 4°C overnight until further processing. The sample was rinsed in 0.1M Cacodylate buffer for 2×5min and postfixed using 1per cent OsO4 in 0.1M Cacodylate buffer, for 1h at 4°C. Then, sample was rinsed for 2×10 min in 0.1M Cacodlyate buffer and secondary post-fixed with 4per cent water solution of uranyl acetate, 1h at RT. Rotation was used at all steps of sample processing. Followed by 5 min wash in MiliQ water, the sample was stepwise dehydrated in Ethanol (25per cent, 50per cent each 15min, 95per cent, 3X100per cent each 20 min) and infiltrated in a graded series of Epon (Ethanol/Epon 3/1, 1/1, 1/3, each 45 min). Sample was left in absolute Epon (EmBed812) overnight. The following day, sample was placed in a fresh absolute Epon for 1h and polymerized (flat embedded) at 60°C for 24-48h. Once polymerized, most surrounding Epon was cut off using razorblade and sample was mounted on empty Epon blocks (samples flat on the top of the blocks) and left at 60 °C for 24h-48h. The region of interest was targeted by ultramicrotome, sections stained with toluidine blue and compared with the MicroCT and LM datasets. After targeting, serial 70nm sections were collected in formvar coated slot grids. The sections were post stained with uranyl acetate (4per cent) and lead citrate. The sections were imaged in a Biotwin CM120 Philips (FEI) TEM at 80kV with a SIS 1K KeenView. Stitches of the 70 sections were aligned using the Track EM plugin in Fiji (Cardona et al., 2012).

### Blood flow mapping

For flow speed mapping, we injected MemGlow labelled 4T1 EVs in Tg (kdrl:EGFP; gata1:DsRed) embryos in Casper background at 48hpf. 30 min post injection, we recorded short time-lapses in 4 different regions in the zebrafish caudal plexus (two in the dorsal aorta and two in the venous area). For each region, we recorded 30s time-lapses (17ms time interval) at single focal plane with flowing red blood cells (RBC). Flow speed was measured by tracking RBC in each region with IMARIS software. tEVs (Cy5) accumulation in endothelial cells (GFP) was measured by IMARIS for each region.

### Pharmacological blood flow tuning

IBMX (3-Isobutyl-1-Methylxanthin – 15879 - Merck) and lidocaine (L7757 – Merck) were directly added to the embryo water (Danieau/PTU solution) containing ZF embryos at the concentration of 100μM in DMSO for 20h and at the concentration of 640μM in EtOH for 2h respectively. Control embryos were treated with similar amount of DMSO or EtOH accordingly. Embryos were then mounted and maintained in fresh solutions of drugs until the end of the experiment. Heartbeat of the embryos were recorded as short time-lapses with a Stereomicroscope (Leica M205 FA). Heartbeats were manually counted on kymographs.

### In vivo Angiogenesis assay

For angiogenesis assay, PTU treated Tg(Fli:LA-eGFP) dechorionated embryos were either uninjected (control) or injected with PBS or with 4T1 Memglow-labeled tEVs at 36 hpf intravascularly. At 48 hpf, PBS injected and tEV injected embryos received 4T1–NLS-tdTomato-IH tumor cell injection (volume-18 nl containing 100-150 cells) in the perivitelline space. 24 hpi of tumor cells embryos were imaged by spinning Disk focusing on the sub-intestinal vein (SIV) plexus (that spans the dorso-lateral part of the yolk) as described in (Hen et al., 2015). The number of newly formed vascular sprouts were manually counted.

### Transcriptomic analyses

Transcriptomics from Fig. 4a was performed on HUVECs cultured in static or flow conditions and treated with 1.109 EVs or PBS for 24h, while transcriptomics from Fig.3a was performed on HUVECs as described previously (Follain et al., 2021). RNA extraction and sequencing Total RNA was extracted using RNeasy Mini Kit (Qiagen, Hilden, Germany) and RNA integrity was assessed by Bioanalyzer (total RNA Pico Kit, 2100 Instrument, Agilent Technologies, Paolo Alto, USA). SMART-Seq^®^ HT PLUS Kit (Takara, Kusatsu, Japan) was used to build mRNA libraries. Libraries were pooled and sequenced (single-end, 75bp) on a NextSeq500 using the NextSeq 500/550 High Output Kit v2 according to the manufacturer’s instructions (Illumina Inc., San Diego, CA, USA). Analysis of RNA-sequence reads: Identification of differentially expressed genes Sequence reads were mapped on the human hg19 genome using STAR (Dobin et al., 2013) to obtain a BAM file (Binary Alignment Map). Raw read counts were determined as an abundance matrix with the HTseq-count tool of the Python package HTSeq (Anders et al., 2015). Trimmed Mean of M-values normalization (TMM) was applied using the EdgeR package (Robinson et al., 2010). A voom transformation was applied to the data that were then fitted into a linear model using weighted least squares for each gene with limma package (Ritchie et al., 2015). Finally, a contrast matrix was created and differential expressions were computed. Up- and down-regulated genes were selected based on adjusted p-values <0.1 and log2 fold-changes > 0.5 and Functional enrichment analyses were performed using STRING v11 (Szklarczyk et al., 2019) and Gene Ontology (Carbon et al., 2021).

### Statistical analyses

All experiments were performed and results were analyzed in at least three independent experiments. Statistical analysis of the results was done using GraphPad Prism (Software version 9.0). Normality of the data was confirmed using Shapiro-Wilkson test and accordingly different statistical tests were used as described in legends. For data that follow gaussian distribution, unpaired t test or one way Anova (with Tukey’s post-test analysis) were used. For data that do not follow gaussian distribution, Mann-Whitney or Kruskal-Wallis tests (with Dunn’s post-test analysis) were used. Illustrations of the statistical analyses were displayed in the figures as the mean +/- standard deviation (SD). p-Values smaller than 0.05 were considered as statistically significant. *, p<0.05, **, p<0.01, ***, p<0.001, ****, p<0.0001.

## Acknowledgements

We thank all members of the Goetz Lab for helpful discussions, in particular Florent Colin for careful reading, Naёl Osmani and Gautier Follain for early interaction on HUVEC analysis. We thank Pascal Kessler (PIC-STRA, CRBS) for assistance in imaging and image analysis, as well as Gregory Khelifi and Camille Hergott for animal care. We are grateful to Nicolas Vitale (INCI, Strasbourg, France), K. C. Kondapalli (Michigan-Dearborn University, USA) and A. M. Weaver (Vanderbilt University, USA) for sharing constructs, and to Yannick Schwab and Pedro Machado (EMBL, Germany) for help with the iCLEM analysis. This work was supported by a fourth-year thesis and a post-doctoral fellowship from la Fondation pour Rercherche Medicale (FRM) to BM and KJR respectively; by grants from La Ligue contre le Cancer, Cancéropôle GrandEst, INCa (PLBIO19-291) and Plan Cancer (3R and Nanotumor) to VH and JGG.; and by institutional funds from University of Strasbourg and INSERM.

## Author Contributions

BM and NA performed most experiments, with help from KJR (image analysis), AL (molecular biology, cell line generation), and OL (molecular biology, cell line generation). Electron microscopy processing (including iCLEM) and analysis was performed by IB and VH. TS, AP, AM and RC performed RNA sequencing and analysis. JGG and VH supervised and designed the project. JGG (with help from VH) was responsible for funding the project. JGG and VH wrote the manuscript with insights from all authors.

---

**Fig. S1.**
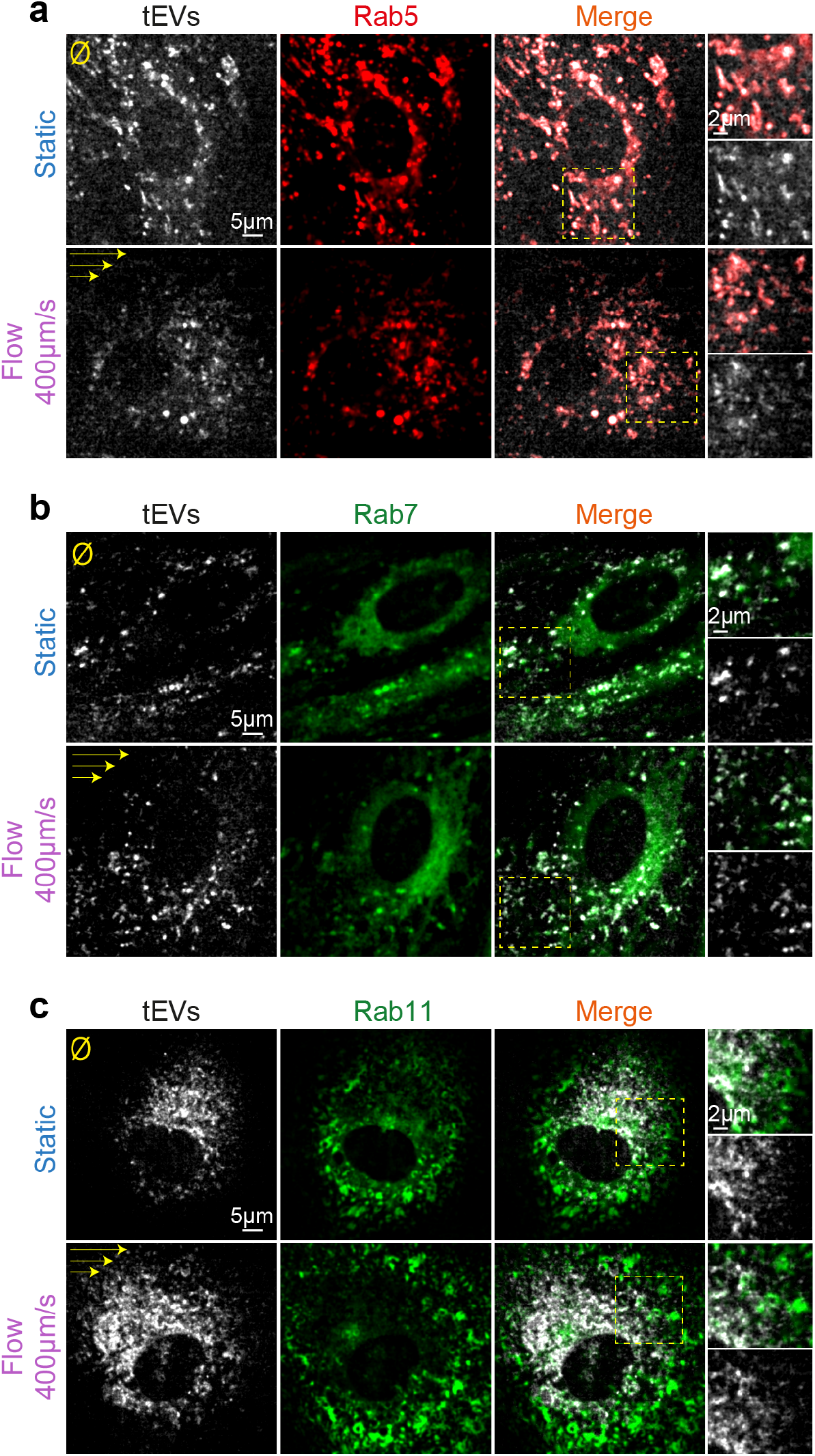
tEVs trafficking in endothelial cells in flow and static conditions. Representative confocal images (single plane) of internalized EVs in vHUVECs cells cultured in flow or static conditions and expressing either mCherry-RAB5 (a), mEmerald-RAB7 (b), eGFP-RAB11 (c).

**Supplementary material** Movie 1: Spinning disk imaging of red blood cells in four different regions of the zebrafish caudal plexus, as described in Figure 1a.

**Fig. S2.**
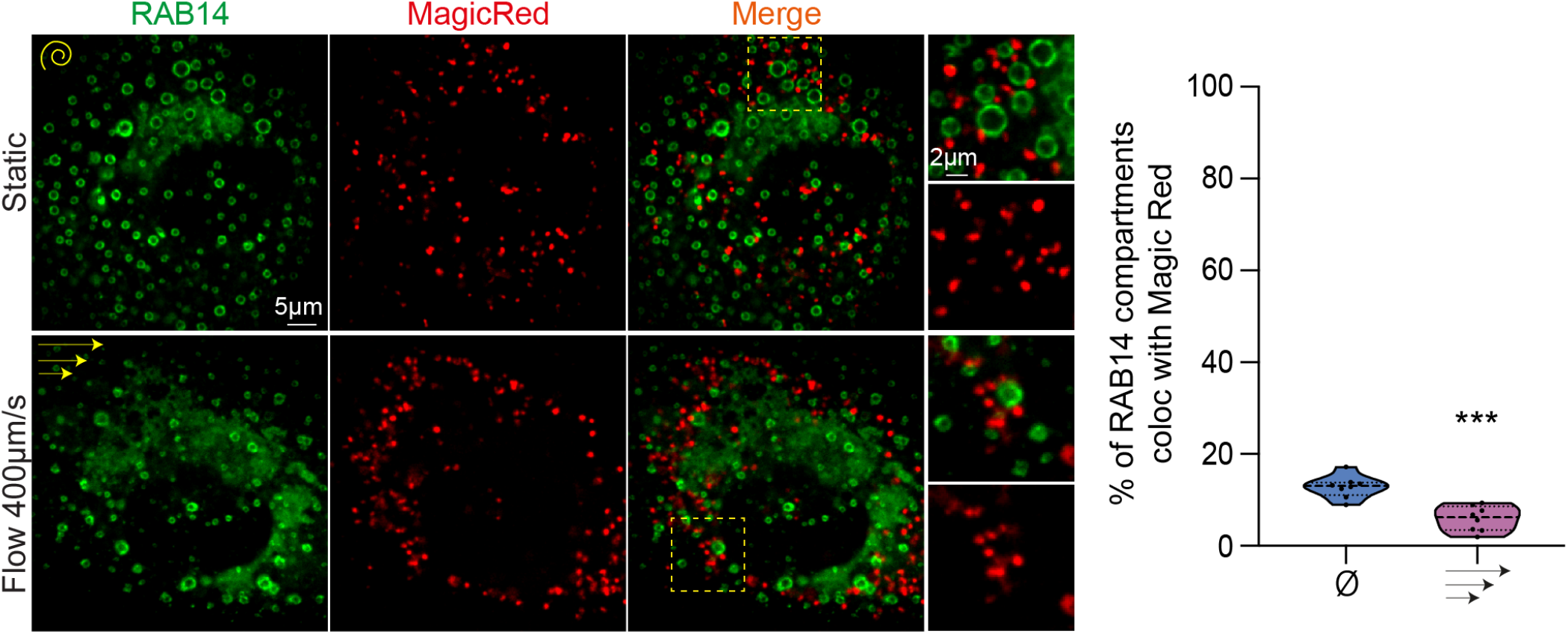
RAB14 compartments show low colocalization with Magic Red in flow and static conditions. Representative confocal images (single plane) of vHUVECs cells expressing GFP-RAB14 cultured in flow or static conditions with Magic Red. (1 dot = 1 FOV p=0,0001 Mann Whitney).

## Bibliography

de Araujo, M.E.G., G. Liebscher, M.W. Hess, and L.A. Huber. 2020. Lysosomal size matters. Traffic. 21:60–75. doi:10.1111/TRA.12714/.

Baron, M. 2012. Endocytic routes to Notch activation. Semin. Cell Dev. Biol. 23:437–442. doi:10.1016/J.SEMCDB.2012.01.008.

Benezra, R., S. Rafii, and D. Lyden. 2001. The Id proteins and angiogenesis. Oncogene. 20:8334–8341. doi:10.1038/sj/onc/1205160.

Beydoun, R., M.A. Hamood, D.M. Gomez Zubieta, and K.C. Kondapalli. 2017. Na+/H+ exchanger 9 regulates iron mobilization at the blood-brain barrier in response to iron starvation. J. Biol. Chem. 292:4293–4301. doi:10.1074/jbc.M116.769240.

Bonsergent, E., E. Grisard, J. Buchrieser, O. Schwartz, C. Théry, and G. Lavieu. 2021. Quantitative characterization of extracellular vesicle uptake and content delivery within mammalian cells. Nat. Commun. 12:1–11. doi:10.1038/s41467-021-22126-y.

Carbon, S., E. Douglass, B.M. Good, D.R. Unni, N.L. Harris, C.J. Mungall, S. Basu, R.L. Chisholm, R.J. Dodson, E. Hartline, P. Fey, P.D. Thomas, L.P. Albou, D. Ebert, M.J. Kesling, H. Mi, A. Muruganujan, X. Huang, T. Mushaya-hama, S.A. LaBonte, D.A. Siegele, G. Antonazzo, H. Attrill, N.H. Brown, P. Garapati, S.J. Marygold, V. Trovisco, G. dos Santos, K. Falls, C. Tabone, P. Zhou, J.L. Goodman, V.B. Strelets, J. Thurmond, P. Garmiri, R. Ishtiaq, M. Rodríguez-López, M.L. Acencio, M. Kuiper, A. Lægreid, C. Logie, R.C. Lovering, B. Kramarz, S.C.C. Saverimuttu, S.M. Pinheiro, H. Gunn, R. Su, K.E. Thurlow, M. Chibucos, M. Giglio, S. Nadendla, J. Munro, R. Jackson, M.J. Duesbury, N. Del-Toro, B.H.M. Meldal, K. Paneerselvam, L. Perfetto, P. Porras, S. Orchard, A. Shrivastava, H.Y. Chang, R.D. Finn, A.L. Mitchell, N.D. Rawlings, L. Richardson, A. Sangrador-Vegas, J.A. Blake, K.R. Christie, M.E. Dolan, H.J. Drabkin, D.P. Hill, L. Ni, D.M. Sitnikov, M.A. Harris, S.G. Oliver, K. Rutherford, V. Wood, J. Hayles, J. Bähler, E.R. Bolton, J.L. de Pons, M.R. Dwinell, G.T. Hayman, M.L. Kaldunski, A.E. Kwitek, S.J.F. Laulederkind, C. Plasterer, M.A. Tutaj, M. Vedi, S.J. Wang, P. D’Eustachio, L. Matthews, J.P. Balhoff, S.A. Aleksander, M.J. Alexander, J.M. Cherry, S.R. Engel, et al. 2021. The Gene Ontology resource: Enriching a GOld mine. Nucleic Acids Res. 49:D325–D334. doi:10.1093/nar/gkaa1113.

Cardona, A., S. Saalfeld, J. Schindelin, I. Arganda-Carreras, S. Preibisch, M. Longair, P. Tomancak, V. Hartenstein, and R.J. Douglas. 2012. TrakEM2 software for neural circuit reconstruction. PLoS One. 7. doi:10.1371/journal.pone.0038011.

De Chaumont, F., S. Dallongeville, N. Chenouard, N. Hervé, S. Pop, T. Provoost, V. Meas-Yedid, P. Pankajakshan, T. Lecomte, Y. Le Montagner, T. Lagache, A. Dufour, and J.C. Olivo-Marin. 2012. Icy: An open bioimage informatics platform for extended reproducible research. Nat. Methods. 9:690–696. doi:10.1038/nmeth.2075.

Chen, P.Y., L. Qin, G. Li, Z. Wang, J.E. Dahlman, J. Malagon-Lopez, S. Gujja, N.A. Cilfone, K.J. Kauffman, L. Sun, H. Sun, X. Zhang, B. Aryal, A. Canfran-Duque, R. Liu, P. Kusters, A. Sehgal, Y. Jiao, D.G. Anderson, J. Gulcher, C. Fernandez-Hernando, E. Lutgens, M.A. Schwartz, J.S. Pober, T.W. Chittenden, G. Tellides, and M. Simons. 2019. Endothelial TGF-β signalling drives vascular inflammation and atherosclerosis. Nat. Metab. 2019 19. 1:912–926. doi:10.1038/S42255-019-0102-3.

Chen, Y.Y., A.M. Syed, P. MacMillan, J. V. Rocheleau, and W.C.W. Chan. 2020. Flow Rate Affects Nanoparticle Uptake into Endothelial Cells. Adv. Mater. 32:1–7. doi:10.1002/adma.201906274.

Cheng, L., and A.F. Hill. 2022. Therapeutically harnessing extracellular vesicles. Nat. Rev. Drug Discov. 2022 215. 21:379–399. doi:10.1038/S41573-022-00410-W.

Collot, M., P. Ashokkumar, H. Anton, E. Boutant, O. Faklaris, T. Galli, Y. Mély, L. Danglot, and A.S. Klymchenko. 2018. MemBright: a family of red to near-infrared fluorescent membrane probes for advanced cellular imaging and neuroscience.

Fang, Y., D. Wu, and K.G. Birukov. 2019. Mechanosensing and Mechanoregulation of Endothelial Cell Functions. Compr. Physiol. 9:873. doi:10.1002/CPHY.C180020.

Fischer, A., N. Schumacher, M. Maier, M. Sendtner, and M. Gessler. 2004. The Notch target genes Hey1 and Hey2 are required for embryonic vascular development. doi:10.1101/gad.291004.

Follain, G., D. Herrmann, S. Harlepp, V. Hyenne, N. Osmani, S.C. Warren, P. Timpson, and J.G. Goetz. 2020. Fluids and their mechanics in tumour transit: shaping metastasis. Nat. Rev. Cancer. 20:107–124. doi:10.1038/s41568-019-0221-x.

Follain, G., N. Osmani, V. Gensbittel, N. Asokan, A. Larnicol, L. Mercier, M.J. Garcia-Leon, I. Busnelli, A. Pichot, N. Paul, R. Carapito, S. Bahram, O. Lefebvre, and J.G. Goetz. 2021. Impairing flow-mediated endothelial remodeling reduces extravasation of tumor cells. Sci. Rep. 11:1–15. doi:10.1038/s41598-021-92515-2.

Follain, G., N. Osmani, A. Sofia, G. Allio, L. Mercier, M.A. Karreman, G. Solecki, M.G. Leon, O. Lefebvre, N. Fekonja, C. Hille, V. Chabannes, G. Dollé, T. Metivet, F. Der Hovsepian, C. Prud’Homme, A. Pichot, N. Paul, R. Carapito, S. Bahram, B. Ruthensteiner, A. Kemmling, S. Siemonsen, T. Schneider, J. Fiehler, M. Glatzel, F. Winkler, Y. Schwab, K. Pantel, S.S. Harlepp, J.G. Goetz, A.S. Azevedo, G. Allio, L. Mercier, M.A. Karreman, G. Solecki, M.J. Garcia Leòn, O. Lefebvre, N. Fekonja, C. Hille, V. Chabannes, G. Dollé, T. Metivet, F. Der Hovsepian, C. Prudhomme, A. Pichot, N. Paul, R. Carapito, S. Bahram, B. Ruthensteiner, A. Kemmling, S. Siemonsen, T. Schneider, J. Fiehler, M. Glatzel, F. Winkler, Y. Schwab, K. Pantel, S.S. Harlepp, and J.G. Goetz. 2018. Hemodynamic forces tune the arrest, adhesion and extravasation of circulating tumor cells. Dev. Cell. 45:33–52.e12. doi:10.1016/j.devcel.2018.02.015.

Freund, J.B., J.G. Goetz, K.L. Hill, and J. Vermot. 2012. Fluid flows and forces in development: Functions, features and biophysical principles. Dev. 139:3063. doi:10.1242/dev.085902.

García-Silva, S., A. Benito-Martín, L. Nogués, A. Hernández-Barranco, M.S. Mazariegos, V. Santos, M. Hergueta-Redondo, P. Ximénez-Embún, R.P. Kataru, A.A. Lopez, C. Merino, S. Sánchez-Redondo, O. Graña-Castro, I. Matei, J.Á. Nicolás-Avila, R. Torres-Ruiz, S. Rodríguez-Perales, L. Martínez, M. Pérez-Martínez, G. Mata, A. Szumera-Ciećkiewicz, I. Kalinowska, A. Saltari, J.M. Martínez-Gómez, S.A. Hogan, H.U. Saragovi, S. Ortega, C. Garcia-Martin, J. Boskovic, M.P. Levesque, P. Rutkowski, A. Hidalgo, J. Muñoz, D. Megías, B.J. Mehrara, D. Lyden, and H. Peinado. 2021. Melanoma-derived small extracellular vesicles induce lymphangiogenesis and metastasis through an NGFR-dependent mechanism. Nat. Cancer 2021 212. 2:1387–1405. doi:10.1038/S43018-021-00272-Y.

Ghoroghi, S., B. Mary, N. Asokan, J.G. Goetz, and V. Hyenne. 2021a. Tumor extracellular vesicles drive metastasis (it,s a long way from home). FASEB BioAdvances. 3:930–943. doi:10.1096/FBA.2021-00079.

Ghoroghi, S., B. Mary, A. Larnicol, N. Asokan, A. Klein, N. Osmani, I. Busnelli, F. Delalande, N. Paul, S. Halary, F. Gros, L. Fouillen, A.-M. Haeberle, C. Royer, C. Spiegelhalter, G. André-Grégoire, V. Mittelheisser, A. Detappe, K. Murphy, P. Timpson, R. Carapito, M. Blot-Chabaud, J. Gavard, C. Carapito, N. Vitale, O. Lefebvre, J.G. Goetz, and V. Hyenne. 2021b. Ral GTPases promote breast cancer metastasis by controlling biogenesis and organ targeting of exosomes. Elife. doi:10.7554/elife.61539.

Han, J., V. V. Shuvaev, P.F. Davies, D.M. Eckmann, S. Muro, and V.R. Muzykantov. 2015. Flow shear stress differentially regulates endothelial uptake of nanocarriers targeted to distinct epitopes of PECAM-1. J. Control. Release. 210:39–47. doi:10.1016/j.jconrel.2015.05.006.

Han, J., B.J. Zern, V. V. Shuvaev, P.F. Davies, S. Muro, and V. Muzykantov. 2012. Acute and chronic shear stress differently regulate endothelial internalization of nanocarriers targeted to platelet-endothelial cell adhesion molecule-1. ACS Nano. 6:8824–8836. doi:10.1021/nn302687n.

Hen, G., J. Nicenboim, O. Mayseless, L. Asaf, M. Shin, G. Busolin, R. Hofi, G. Almog, N. Tiso, N.D. Lawson, and K. Yaniv. 2015. Venous-derived angioblasts generate organ-specific vessels during zebrafish embryonic development. doi:10.1242/dev.129247.

Hoffman, H.K., R.S. Aguilar, A.R. Clark, N.S. Groves, N. Pezeshkian, M.M. Bruns, and S.B. van Engelenburg. 2022. Endocytosed HIV-1 Envelope Glycoprotein Traffics to Rab14 + Late Endosomes and Lysosomes to Regulate Surface Levels in T-Cell Lines. J. Virol. 96. doi:10.1128/JVI.00767-22.

Hoshino, A., B. Costa-Silva, T.-L. Shen, G. Rodrigues, A. Hashimoto, M. Tesic Mark, H. Molina, S. Kohsaka, A. Di Giannatale, S. Ceder, S. Singh, C. Williams, N. Soplop, K. Uryu, L. Pharmer, T. King, L. Bojmar, A.E. Davies, Y. Ararso, T. Zhang, H. Zhang, J. Hernandez, J.M. Weiss, V.D. Dumont-Cole, K. Kramer, L.H. Wexler, A. Narendran, G.K. Schwartz, J.H. Healey, P. Sandstrom, K. Jørgen Labori, E.H. Kure, P.M. Grandgenett, M.A. Hollingsworth, M. de Sousa, S. Kaur, M. Jain, K. Mallya, S.K. Batra, W.R. Jarnagin, M.S. Brady, O. Fodstad, V. Muller, K. Pantel, A.J. Minn, M.J. Bissell, B.A. Garcia, Y. Kang, V.K. Rajasekhar, C.M. Ghajar, I. Matei, H. Peinado, J. Bromberg, and D. Lyden. 2015. Tumour exosome integrins determine organotropic metastasis. Nature. 1–19. doi:10.1038/nature15756.

Hyenne, V., A. Apaydin, D. Rodriguez, C. Spiegelhalter, S. Hoff-Yoessle, M. Diem, S. Tak, O. Lefebvre, Y. Schwab, J.G. Goetz, and M. Labouesse. 2015. RAL-1 controls multivesicular body biogenesis and exosome secretion. J. Cell Biol. 211:27–37. doi:10.1083/jcb.201504136.

Hyenne, V., S. Ghoroghi, M. Collot, J. Bons, G. Follain, S. Harlepp, B. Mary, J. Bauer, L. Mercier, I. Busnelli, O. Lefebvre, N. Fekonja, M.J. Garcia-Leon, P. Machado, F. Delalande, A.A. López, S.G. Silva, F.J. Verweij, G. van Niel, F. Djouad, H. Peinado, C. Carapito, A.S. Klymchenko, and J.G. Goetz. 2019. Studying the Fate of Tumor Extracellular Vesicles at High Spatiotemporal Resolution Using the Zebrafish Embryo. Dev. Cell. 48:554–572.e7. doi:10.1016/j.devcel.2019.01.014.

Imai, T., Y. Takahashi, M. Nishikawa, K. Kato, M. Morishita, T. Yamashita, A. Matsumoto, C. Charoenviriyakul, and Y. Takakura. 2015. Macrophage-dependent clearance of systemically administered B16BL6-derived exosomes from the blood circulation in mice. J. Extracell. vesicles. 4:26238. doi:10.3402/jev.v4.26238.

Jerabkova-Roda, K., A. Dupas, N. Osmani, V. Hyenne, and J. G. Goetz. 2022. Circulating extracellular vesicles and tumor cells: sticky partners in metastasis. Trends in Cancer. 8:799–805. doi:10.1016/J.TRECAN.2022.05.002.

Joshi, B.S., M.A. de Beer, B.N.G. Giepmans, and I.S. Zuhorn. 2020. Endocytosis of Extracellular Vesicles and Release of Their Cargo from Endosomes. ACS Nano. doi:10.1021/acsnano.9b10033.

Kalluri, R., and V.S. LeBleu. 2020. The biology, function, and biomedical applications of exosomes. Science (80-.). 367. doi:10.1126/science.aau6977.

Kienast, Y., L. von Baumgarten, M. Fuhrmann, W.E.F. Klinkert, R. Goldbrunner, J. Herms, and F. Winkler. 2010. Realtime imaging reveals the single steps of brain metastasis formation. Nat. Med. 16:116–122. doi:10.1038/nm.2072.

Kitagawa, M., M. Hojo, I. Imayoshi, M. Goto, M. Ando, T. Ohtsuka, R. Kageyama, and S. Miyamoto. 2013. Hes1 and Hes5 regulate vascular remodeling and arterial specification of endothelial cells in brain vascular development. Mech. Dev. 130:458–466. doi:10.1016/J.MOD.2013.07.001.

Kondapalli, K.C., J.P. Llongueras, V. Capilla-González, H. Prasad, A. Hack, C. Smith, H. Guerrero-Cázares, A. Quiñones-Hinojosa, and R. Rao. 2015. A leak pathway for luminal protons in endosomes drives oncogenic signalling in glioblastoma. Nat. Commun. 6. doi:10.1038/ncomms7289.

Kuijl, C., M. Pilli, S.K. Alahari, H. Janssen, P.S. Khoo, K. E. Ervin, M. Calero, S. Jonnalagadda, R.H. Scheller, J. Neefjes, and J.R. Junutula. 2013. Rac and Rab GTPases dual effector Nischarin regulates vesicle maturation to facilitate survival of intracellular bacteria. EMBO J. 32:713–727. doi:10.1038/EMBOJ.2013.10.

Kyei, G.B., I. Vergne, J. Chua, E. Roberts, J. Harris, J.R. Junutula, and V. Deretic. 2006. Rab14 is critical for maintenance of Mycobacterium tuberculosis phagosome maturation arrest. EMBO J. 25:5250–5259. doi:10.1038/SJ.EMBOJ.7601407.

Li, Y.S.J., J.H. Haga, and S. Chien. 2005. Molecular basis of the effects of shear stress on vascular endothelial cells. J. Biomech. 38:1949–1971. doi:10.1016/J.JBIOMECH.2004.09.030.

Lin, A., A. Sabnis, S. Kona, S. Nattama, H. Patel, J.F. Dong, and K.T. Nguyen. 2010. Shear-regulated uptake of nanoparticles by endothelial cells and development of endothelial-targeting nanoparticles. J. Biomed. Mater. Res. Part A. 93A:833–842. doi:10.1002/JBM.A.32592.

Marar, C., B. Starich, and D. Wirtz. 2021. Extracellular vesicles in immunomodulation and tumor progression. Nat. Immunol. 22:560–570. doi:10.1038/s41590-021-00899-0.

Mary, B., S. Ghoroghi, V. Hyenne, and J.G. Goetz. 2020. Live tracking of extracellular vesicles in larval zebrafish. In Methods in Enzymology.

Morad, G., C. V. Carman, E.J. Hagedorn, J.R. Perlin, L.I. Zon, N. Mustafaoglu, T.E. Park, D.E. Ingber, C.C. Daisy, and M.A. Moses. 2019. Tumor-Derived Extracellular Vesicles Breach the Intact Blood-Brain Barrier via Transcytosis. ACS Nano. 13:13853–13865. doi:10.1021/acsnano.9b04397.

Morioka, T., M. Sakabe, T. Ioka, T. Iguchi, K. Mizuta, M. Hattammaru, C. Sakai, M. Itoh, G.E. Sato, A. Hashimoto, M. Fujita, K. Okumura, M. Araki, M. Xin, R.A. Pedersen, M.F. Utset, H. Kimura, and O. Nakagawa. 2014. An Important Role of Endothelial Hairy-Related Transcription Factors in Mouse Vascular Development. doi:10.1002/dvg.22825.

Morishita, M., Y. Takahashi, M. Nishikawa, K. Sano, K. Kato, T. Yamashita, T. Imai, H. Saji, and Y. Takakura. 2015. Quantitative analysis of tissue distribution of the B16BL6-derived exosomes using a streptavidin-lactadherin fusion protein and Iodine-125-Labeled biotin derivative after intravenous injection in mice. J. Pharm. Sci. 104:705–713. doi:10.1002/jps.24251.

Nicoli, S., and M. Presta. 2007. The zebrafish/tumor xenograft angiogenesis assay. Nat. Protoc. 2:2918–2923. doi:10.1038/nprot.2007.412.

van Niel, G., D.R.F. Carter, A. Clayton, D.W. Lambert, G. Raposo, and P. Vader. 2022. Challenges and directions in studying cell-cell communication by extracellular vesicles. Nat. Rev. Mol. Cell Biol. 23:369–382. doi:10.1038/S41580-022-00460-3.

Okai, B., N. Lyall, N.A.R. Gow, J.M. Bain, and L.P. Erwig. 2015. Rab14 regulates maturation of macrophage phagosomes containing the fungal pathogen Candida albicans and outcome of the host-pathogen interaction. Infect. Immun. 83:1523–1535. doi:10.1128/IAI.02917-14.

Peinado, H., H. Zhang, I.R. Matei, B. Costa-Silva, A. Hoshino, G. Rodrigues, B. Psaila, R.N. Kaplan, J.F. Bromberg, Y. Kang, M.J. Bissell, T.R. Cox, A.J. Giaccia, J.T. Erler, S. Hiratsuka, C.M. Ghajar, and D. Lyden. 2017. Premetastatic niches: organ-specific homes for metastases. Nat. Rev. Cancer. 17:302–317. doi:10.1038/nrc.2017.6.

Perera, R.M., and R. Zoncu. 2016. The Lysosome as a Regulatory Hub. Annu. Rev. Cell Dev. Biol. 32:223–253. doi:10.1146/annurev-cellbio-111315-125125.

Schindelin, J., I. Arganda-Carreras, E. Frise, V. Kaynig, M. Longair, T. Pietzsch, S. Preibisch, C. Rueden, S. Saalfeld, B. Schmid, J.Y. Tinevez, D.J. White, V. Hartenstein, K. Eliceiri, P. Tomancak, and A. Cardona. 2012. Fiji: An open-source platform for biological-image analysis. Nat. Methods. 9:676–682. doi:10.1038/nmeth.2019.

Sheldon, H., E. Heikamp, H. Turley, R. Dragovic, P. Thomas, C. E. Oon, R. Leek, M. Edelmann, B. Kessler, R.C.A. Sainson, I. Sargent, J.L. Li, and A.L. Harris. 2010. New mechanism for Notch signaling to endothelium at a distance by Delta-like 4 incorporation into exosomes. Blood. 116:2385–2394. doi:10.1182/BLOOD-2009-08-239228.

Shelke, G.V., Y. Yin, S.C. Jang, C. Lässer, S. Wennmalm, H.J. Hoffmann, L. Li, Y.S. Gho, J.A. Nilsson, and J. Lötvall. 2019. Endosomal signalling via exosome surface TGFβ-1. J. Extracell. Vesicles. 8. doi:10.1080/20013078.2019.1650458/SUPPL_*F*_ILE/ZJEV_*A*1_650458_*S*_M9071.ZIP.

Stirling, D.R., M.J. Swain-Bowden, A.M. Lucas, A.E. Carpenter, B.A. Cimini, and A. Goodman. 2021. CellProfiler 4: improvements in speed, utility and usability. BMC Bioinformatics. 22:1–11. doi:10.1186/S12859-021-04344-9/FIGURES/6.

Sung, B.H., A. von Lersner, J. Guerrero, E.S. Krystofiak, D. Inman, R. Pelletier, A. Zijlstra, S.M. Ponik, and A.M. Weaver. 2020. A live cell reporter of exosome secretion and uptake reveals pathfinding behavior of migrating cells. Nat. Commun. 11:1–15. doi:10.1038/s41467-020-15747-2.

Szklarczyk, D., A.L. Gable, D. Lyon, A. Junge, S. Wyder, J. Huerta-Cepas, M. Simonovic, N.T. Doncheva, J.H. Morris, P. Bork, L.J. Jensen, and C. Von Mering. 2019. STRING v11: Protein-protein association networks with increased coverage, supporting functional discovery in genome-wide experimental datasets. Nucleic Acids Res. doi:10.1093/nar/gky1131.

Takahashi, Y., M. Nishikawa, H. Shinotsuka, Y. Matsui, S. Ohara, T. Imai, and Y. Takakura. 2013. Visualization and in vivo tracking of the exosomes of murine melanoma B16-BL6 cells in mice after intravenous injection. J. Biotechnol. 165:77–84. doi:10.1016/j.jbiotec.2013.03.013.

Tarbell, J.M. 2010. Shear stress and the endothelial transport barrier. Cardiovasc. Res. 87:320. doi:10.1093/CVR/CVQ146.

Tkach, M., and C. Théry. 2016. Communication by Extracellular Vesicles: where we are and where to go. doi:10.1016/j.cell.2016.01.043.

Todorova, D., S. Simoncini, R. Lacroix, F. Sabatier, and F. Dignat-George. 2017. Extracellular Vesicles in Angiogenesis. Circ. Res. 120:1658–1673. doi:10.1161/CIRCRESAHA.117.309681.

Trofimenko, E., Y. Homma, M. Fukuda, and C. Widmann. 2021. The endocytic pathway taken by cationic substances requires Rab14 but not Rab5 and Rab7. Cell Rep. 37. doi:10.1016/J.CELREP.2021.109945.

Verweij, F.J., L. Balaj, C.M. Boulanger, D.R.F. Carter, E.B. Compeer, G. D’Angelo, S. El Andaloussi, J.G. Goetz, J.C. Gross, V. Hyenne, E.M. Krämer-Albers, C.P. Lai, X. Loyer, A. Marki, S. Momma, E.N.M. Nolte-‘t Hoen, D.M. Pegtel, H. Peinado, G. Raposo, K. Rilla, H. Tahara, C. Théry, M.E. van Royen, R.E. Vandenbroucke, A.M. Wehman, K. Witwep Z. Wu, R. Wubbolts, and G. van Niel. 2021. The power of imaging to understand extracellular vesicle biology in vivo. Nat. Methods. 18:1013–1026. doi:10.1038/S41592-021-01206-3.

Verweij, F.J., V. Hyenne, G. Van Niel, and J.G. Goetz. 2019a. Extracellular Vesicles: Catching the Light in Zebrafish. Trends Cell Biol. 29:770–776. doi:10.1016/j.tcb.2019.07.007.

Verweij, F.J., C. Revenu, G. Arras, F. Dingli, D. Loew, D.M. Pegtel, G. Follain, G. Allio, J.G. Goetz, P. Zimmermann, P. Herbomel, F. Del Bene, G. Raposo, and G. van Niel. 2019b. Live Tracking of Inter-organ Communication by Endogenous Exosomes In Vivo. Dev. Cell. 48:573–589.e4. doi:10.1016/j.devcel.2019.01.004.

Vion, A.C., M. Kheloufi, A. Hammoutene, J. Poisson, J. Lasselin, C. Devue, I. Pic, N. Dupont, J. Busse, K. Stark, J. Lafaurie-Janvore, A.I. Barakat, X. Loyer, M. Souyri, B. Viollet, P. Julia, A. Tedgui, P. Codogno, C.M. Boulanger, and P.E. Rautou. 2017. Autophagy is required for endothelial cell alignment and atheroprotection under physiological blood flow. Proc. Natl. Acad. Sci. U. S. A. doi:10.1073/pnas.1702223114.

Wieland, E., J. Rodriguez-Vita, S.S. Liebler, C. Mogler, I. Moll, S.E. Herberich, E. Espinet, E. Herpel, A. Menuchin, J. Chang-Claude, M. Hoffmeister, C. Gebhardt, H. Brenner, A. Trumpp, C.W. Siebel, M. Hecker, J. Utikal, D. Sprinzak, and A. Fischer. 2017. Endothelial Notch1 Activity Facilitates Metastasis. Cancer Cell. 31:355–367. doi:10.1016/J.CCELL.2017.01.007.

Xie, F., X. Zhou, P. Su, H. Li, Y. Tu, J. Du, C. Pan, X. Wei, M. Zheng, K. Jin, L. Miao, C. Wang, X. Meng, H. van Dam, P. ten Dijke, L. Zhang, and F. Zhou. 2022. Breast cancer cell-derived extracellular vesicles promote CD8+ T cell exhaustion via TGF-β type II receptor signaling. Nat. Commun. 13. doi:10.1038/s41467-022-31250-2.

Yáñez-Mó, M., P.R.-M. Siljander, Z. Andreu, A.B. Zavec, F.E. Borràs, E.I. Buzas, K. Buzas, E. Casal, F. Cappello, J. Carvalho, E. Colás, A. Cordeiro-da Silva, S. Fais, J.M. Falcon-Perez, I.M. Ghobrial, B. Giebel, M. Gimona, M. Graner, I. Gursel, M. Gursel, N.H.H. Heegaard, A. Hendrix, P. Kierulf, K. Kokubun, M. Kosanovic, V. Kralj-Iglic, E.-M. Krämer-Albers, S. Laitinen, C. Lässer, T. Lener, E. Ligeti, A. Linē, G. Lipps, A. Llorente, J. Lötvall, M. Manček-Keber, A. Marcilla, M. Mittelbrunn, I. Nazarenko, E.N.M. Nolte-‘t Hoen, T.A. Nyman, L. O’Driscoll, M. Olivan, C. Oliveira, É. Pállinger, H.A. Del Portillo, J. Reventós, M. Rigau, E. Rohde, M. Sammar, F. Sánchez-Madrid, N. Santarém, K. Schallmoser, M.S. Ostenfeld, W. Stoorvogel, R. Stukelj, S.G. Van der Grein, M.H. Vasconcelos, M.H.M. Wauben, and O. De Wever. 2015. Biological properties of extracellular vesicles and their physiological functions. J. Extracell. vesicles. 4:27066. doi:10.3402/jev.v4.27066.

